# Various tomato cultivars display contrasted morphological and molecular responses to a heat wave

**DOI:** 10.1101/2022.09.08.507118

**Authors:** N. Bollier, R. Micol-Ponce, A. Dakdaki, E. Maza, M. Zouine, A. Djari, M. Bouzayen, C. Chevalier, F. Delmas, N. Gonzalez, M. Hernould

## Abstract

Climate change is one of the biggest threats that human society currently needs to face. Heat waves associated with global warming negatively affect plant growth and development and will increase in frequency. Tomato is one of the most produced and consumed fruit in the world but remarkable yield losses occur every year due to the sensitivity of many cultivars to heat stress. New insights into how tomato plants are responding to heat waves will contribute to the development of new cultivars with high yields under harsh temperature conditions. In this study, the analysis of microsporogenesis and pollen germination rate of eleven tomato cultivars after exposure to a simulated heat wave revealed differences between genotypes. The transcriptome of floral buds at two developmental stages of five cultivars selected based on their pollen germination tolerance or sensitivity, revealed common and specific molecular responses implemented by tomato cultivars to cope with heat waves. These data provide valuable insights into the underlying molecular adaptation of floral buds to heat stress and will contribute to the development of future climate resilient tomato varieties.

## Introduction

Population growth and global warming are two of the major issues that humanity will have to face in the next decades (IPCC, 2014). Based on current climatic models, experts commonly anticipate that rising temperatures will cause substantial yield losses in the future affecting major plant-food sources (Tripathi *et al*., 2016). Global warming has already a significant impact on plant crop production and agricultural practices. During the past recent years, many regions of the world experienced exceptional extreme heat waves causing severe damages to ecosystems, human society and crop production (Stillman, 2019). The frequency and intensity of these heat waves will increase in many regions of the globe leading to major losses in agricultural yield (Lau and Nath, 2012; Molina *et al*., 2020).

Plant growth and development can be strongly altered by high temperatures, the most heat-susceptible phases being early seedling and reproductive phases. Heat-stress (HS) can impair essential cellular, physiological and developmental processes, including genome integrity, photosynthesis, flower development or pollen development and viability as reported for a number of plant species (Han *et al*., 2021; Moore *et al*., 2021). Detrimental effects of heat on pollen production have been described in various plant taxa, including *Arabidopsis thaliana* (Arabidopsis), *Solanum lycopersicum* (tomato) and *Oryza sativa* (rice) and for different developmental stages such as meiosis, microspore development and pollen maturation (Raja *et al*., 2019). Pollen development is known to be one of the most temperature-sensitive processes in all plant life cycle (Zinn *et al*., 2010), and impaired pollen development results in poor fertilization and reduced fruit and seed yield (Giorno *et al*., 2013; Müller and Rieu, 2016). The cellular responses to HS have been studied in a variety of plant species and various organ and tissues revealing both common and specific responses. HS disturbs various cellular processes, among which the denaturation of biological molecules, such as proteins, lipids, nucleic acids (Bohnert *et al*., 2006; Kotak *et al*., 2007, Essemine *et al*., 2010) and the disintegration of subcellular structures, including membranes and cytoskeleton networks (Savchenko *et al*., 2011; Saidi *et al*., 2011) have been described. HS affects genome integrity by triggering DNA damage through nucleotide modifications and single-strand or double-strand breaks (Kantidze *et al*., 2016) and it also alters chromatin architecture (Pecinka *et al*., 2012), and affects meiotic recombination (Ning *et al*., 2021; De Jaeger-Brat *et al*., 2021). To cope with HS, plants have developed highly complex intracellular signaling systems involving hormones, Ca2+ and reactive oxygen species (ROS) (Wahid *et al*., 2007). For a long time, ROS were considered as a byproduct that impairs plant growth. However, in recent period, ROS gained attention for its function as a signaling molecule involved in response to environmental stresses like HS (Medina *et al*. 2021). In support to this idea, it has been shown that ROS-scavenging enzymes, involved in detoxification processes, are rapidly induced by HS (Suzuki and Mittler, 2006).

HS induces rapid transcriptional changes and the elucidation of the complex transcriptional regulatory networks involved in plant responses to HS is now well advanced (Ohama *et al*., 2017). That is, HS rapidly activates HS-responsive transcription factors, HSFs, which regulate the transcription of a wide range of target genes involved in signaling and metabolic pathways (Koskull-Döring *et al*., 2007). Among the most notable responses, the expression of Heat Shock Proteins (HSPs) is known to play essential roles in cellular protection through their action on protein misfolding or aggregation but also in protein translocation and degradation (Vierling 1991; Wang et al. 2004; Schleiff & Becker 2011). Transcriptomic profiling of HS responses in plants has identified several molecular pathways induced under elevated temperature conditions including photosynthesis, response to light or rRNA processing (Jayakodi, Lee and Yang, 2019). Interestingly, in addition to the classically induced heat shock proteins, these studies suggested potential roles for several factors involved in epigenetic, post-transcriptional and post-translational regulation as well as in hormonal regulation. Deciphering the effects of HS at the transcriptome level greatly helped defining candidate genes potentially involved in mediating plant tolerance to HS and several attempts aiming to improve thermotolerance by knocking out or overexpressing these candidate genes have been reported in different plant species including Arabidopsis, tomato, rice and *Glycine max* (soybean) (Zhu *et al*., 2006; Yokotani *et al*., 2008; Xin *et al*., 2010; Wu *et al*., 2012; Li *et al*., 2013; Shen *et al*., 2015; Xue, Drenth and McIntyre, 2015; Wan *et al*., 2016). It is however likely that the activation of two or more independent - but mutually complementary-pathways are able to improve more effectively thermotolerance in crops species. In this regard, new pathways need to be discovered in order to pave the way towards the generation of highly tolerant genotypes using the newly identified candidates for gene stacking.

HS response has been extensively investigated in plants in the past two decades. Most studies have been carried out applying very high temperatures (45°C to 50°C) for a short period of time (from 30 minutes to 3 hours) on the whole plant, or focused on specific developmental stages or organs (Qu *et al*., 2013; Driedonks *et al*., 2016; Janni *et al*., 2020). Strikingly so far, quite a few studies have been carried out that simulate heat wave conditions as defined by the STAtistical and Regional dynamical Downscaling of EXtremes for European regions (Stardex) project: 5 consecutive days with 5 degrees anomaly with respect to mean temperature in summer (Jagadish *et al*., 2021). Because the effect of short-term heat treatments is unlikely to completely reproduce what happens during heat waves, studying the impact of long periods of heat on plants is essential to eventually select plant cultivars with increased heat tolerance.

Tomato is one of the most produced and consumed fruits in the world. However, tomato producers face dramatic yield losses in very warm summers, due to poor fruit setting that leads to lower fruit number and small fruits of low quality (Adams *et al*., 2001). In recent years, several transcriptomic, metabolomic, proteomic or lipidomic analyses were performed to study the response of tomato plants to HS (Pressman *et al*., 2002; Spicher *et al*., 2016; Paupière *et al*., 2017; Almeida *et al*., 2021). Studies focusing on floral buds revealed that short-term HS induces the expression of *HSF* genes, *HSP* genes, ROS scavenger genes and genes involved in the control of sugar levels (Frank *et al*., 2009). However, most of these studies are limited to one tomato genotype and, more importantly, the short-term treatment is unlikely to completely reproduce the molecular responses to heat wave.

Here, we aimed at identifying the common and/or specific molecular responses to a simulated heat wave in a greenhouse, in tomato floral buds of various tomato cultivars. Seeds from 11 modern tomato cultivars were obtained from different seed companies, and plants were grown and subjected to a heat wave, in order to identify tolerant and sensitive cultivars based on pollen germination. Among these 11 cultivars, 2 were selected as thermotolerant and 3 as thermosensitive for subsequent genome-wide transcriptomic profiling analysis of their floral buds, to identify the molecular pathways regulated during heat wave. These analyses highlighted the different strategies that tomato cultivars set-up for facing heat waves.

## Results

### Morphological analyses of flower development under elevated ambient temperatures

To assess phenotypic differences in response to HS of various tomato genotypes, we examined the behavior of eleven tomato cultivars corresponding to ten commercial varieties and the cherry tomato West Virginia 106 (WVA106) cultivar. This set of cultivars was selected in order to cover some of the tomato diversity in terms of i) fruit morphology: small- (WVA106, Sassari, Brioso, M82) medium- (Docet, Moneymaker, Clodano, JAG8810), and large size (Rebelski, Marbonne, DRK7024) (**Figure S1**); ii) growth pattern: determinate (JAG8810, M82) *versus* indeterminate plants (DRK7024, WVA106, Brioso, Marbonne, Sassari, Moneymaker, Rebelski); and **iii)** market suitability: fresh market- (Sassari, Brioso, M82, Clodano, Moneymaker) and processing tomatoes (JAG8810, DOCET). Plants were grown in a greenhouse under non-stress (NS) temperature conditions (24°C during the day and 18°C during the night) until the plants reach the bolting stage. Thereafter, a simulated heat wave was applied, consisting in 3 weeks at 35°C during the day and 25°C during the night. Given that floral development is critical for tomato yield, we focused our analysis on the morphological and cytological changes occurring during this process under elevated ambient temperatures treatment.

Under non-stress (NS) conditions, the 11 selected cultivars displayed variability in size and morphology of floral buds during development (**Figure 1**). Under HS, the size and shape of floral buds were unevenly affected in different cultivars. In most cultivars, floral bud size distribution was similar under NS and HS conditions (**Figure S2**), but clear morphological alterations were observed (**Figure 1**). In particular, the flowers of the Sassari cultivar exhibited recessed stigmas with complete absence of stigma exertion under NS conditions, whereas many flower buds in this cultivar showed stigma exertion under HS. On the other hand, HS induced the appearance of curled petals in Clodano, Docet, M82, Marbonne and Moneymaker cultivars. In addition, HS caused the sepals and petals to open earlier than in NS conditions in Moneymaker, Brioso and WVA106. In this latter cultivar, sepals displayed burned extremities likely due to HS. The number of floral organs developed by all the cultivars studied was the same in both conditions suggesting that HS did not affect floral architecture. In conclusion, floral development in various tomato cultivars was affected differently under HS, suggesting that the response to HS is at least partially dependent on the genetic background.

**Figure 1.**
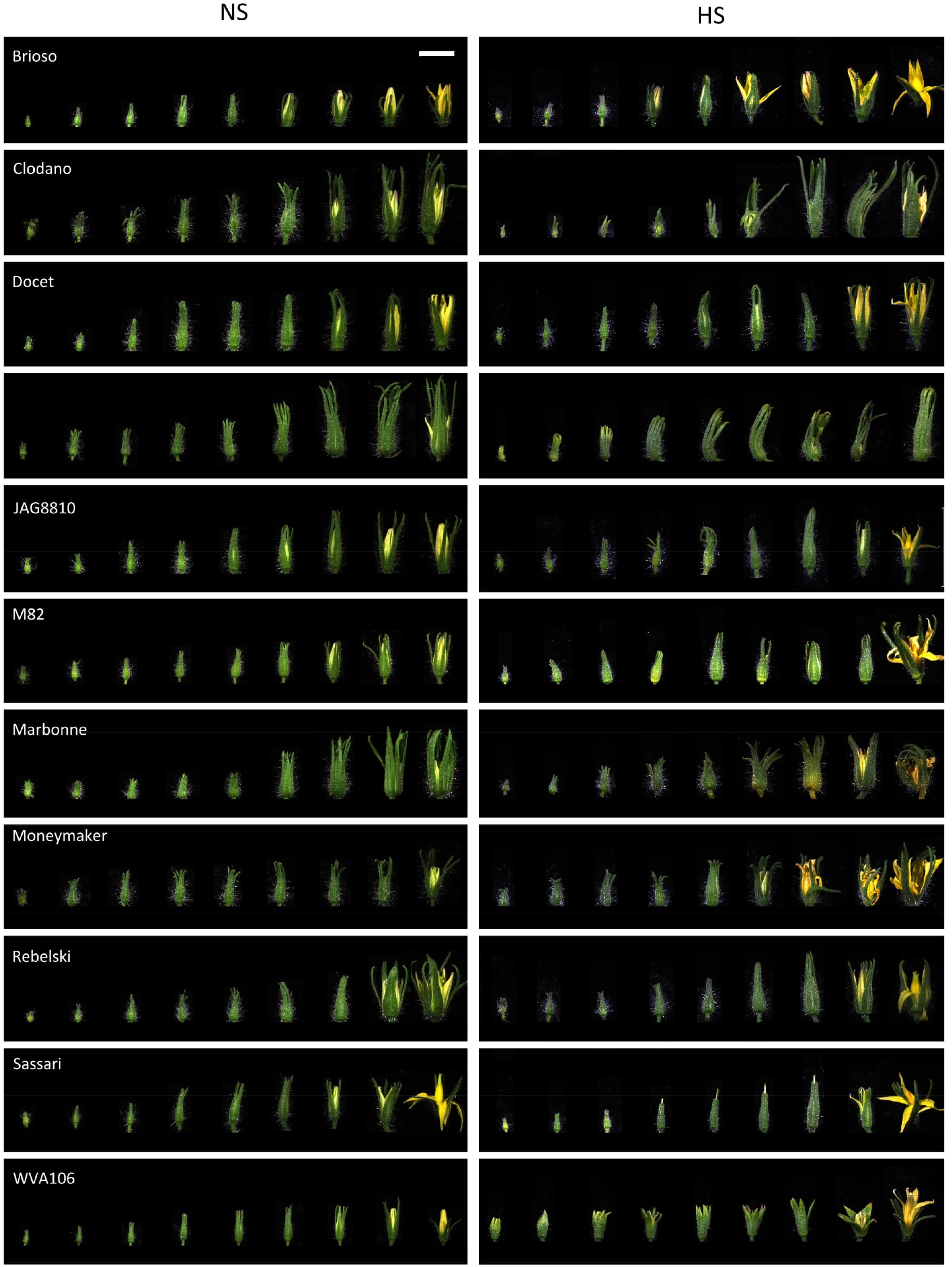
Effects of heat stress (HS) and non-stress (NS) conditions on flower bud growth in the 11 selected tomato cultivars. For each cultivar, floral buds are ordered according to their size in each condition showing the effect of the heat stress at different developmental stages. Scale bar = 1 cm (applies to all panels).

### Variability in pollen germination of tomato cultivars under elevated ambient temperatures

Flower fertilization, and subsequent fruit set and seed production, can be impaired by poor pollen germination and pollen tube growth. In heat-sensitive tomato cultivars, high temperatures reduce pollen production, viability and germination capacity of pollen grains, leading to impaired fruit set and to lower number of seeds per fruit (Firon *et al*., 2006). To investigate the tolerance/sensitivity of pollen germination to HS in the eleven selected tomato cultivars, we determined the percentage of pollen germination as a good proxy of pollen potential for fertilization under NS and HS conditions (**Figure 2**). Under NS conditions, pollen germination was highly variable between the cultivars studied, with a maximum percentage of of 80% for WVA106, and a minimum of 18.6% for DRK7024. Such a variability between tomato cultivars has already been reported and can be attributed to differences of pollen germination capacity between genotypes (Gentile *et al*., 1971; Huner and Huystee, 1982; Abdul-Baki and Stommel, 1995). In addition, this variability can also be due to the composition of the germination medium used which is more or less suitable for some cultivars than others (Karapanos *et al*., 2006). Overall, HS impacted pollen germination negatively in all cultivars, and it prevented even totally any germination for Clodano, DRK7024, Moneymaker, Rebelski and Sassari, indicating the extreme sensitivity to HS of these cultivars (**Figure 2**). In Brioso, Docet, M82 and Marbonne cultivars, the pollen germination rate under HS was reduced respectively by 10-, 21-, 12- and 33-fold, when compared to NS condition, indicating a high sensitivity to HS of these cultivars even though some germination capacity was maintained. Conversely, pollen germination in JAG8810 and WVA106 was only reduced by 1.4- and 3.5-fold respectively, suggesting a better tolerance to HS treatment in these cultivars. Taken together, these results indicated that pollen germination percentage is highly affected under HS conditions although the degree of sensitivity is clearly genotype dependent.

**Figure 2.**
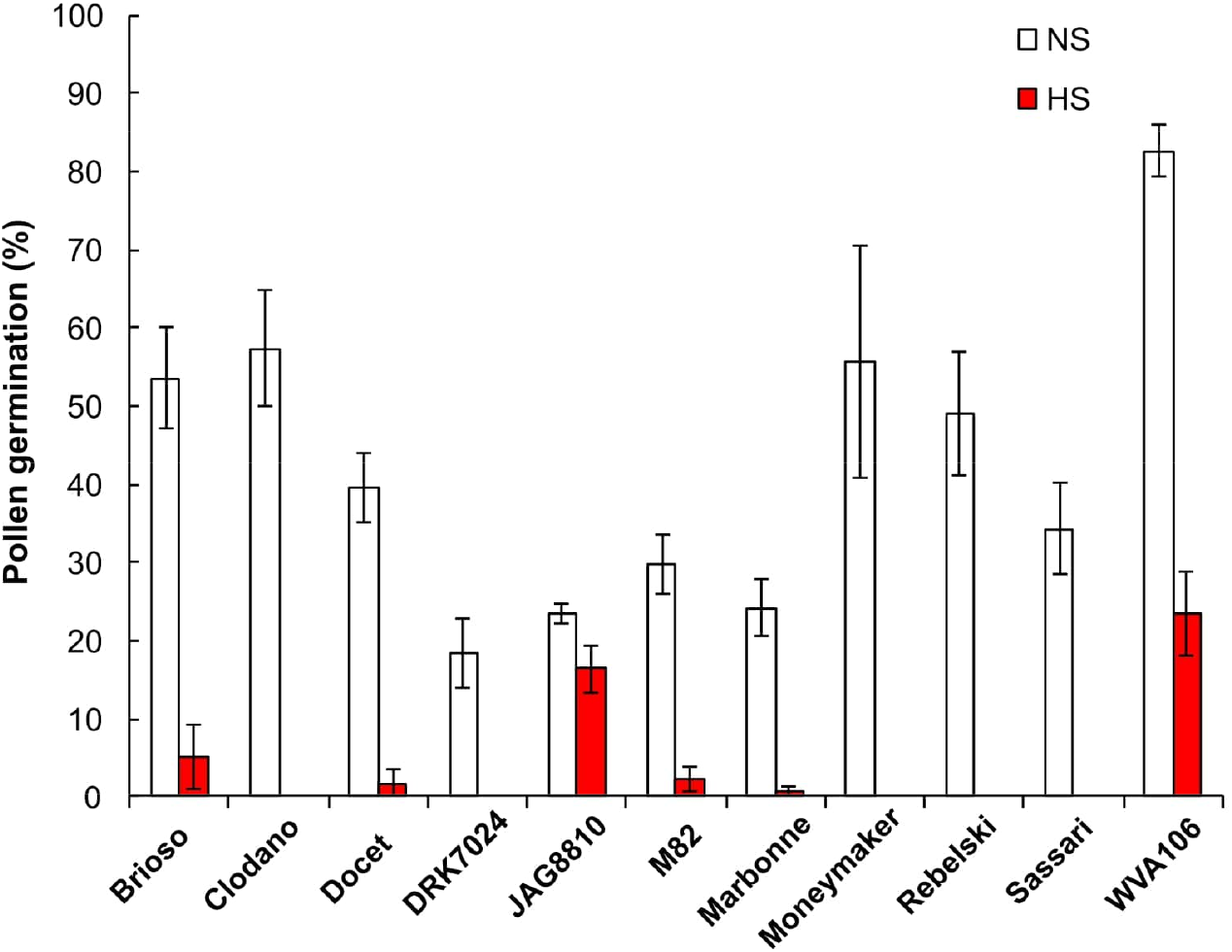
Pollen germination rate expressed as the percentage of germinating pollen grains (%) under non-stress (NS) and heat stress (HS) conditions in 11 tomato cultivars.

### Cytological analyses of pollen development under elevated ambient temperatures

To determine if the effect of the heat wave on pollen germination was related to a perturbation of pollen development, we selected 5 cultivars showing contrasting behaviors for pollen germination under HS: namely WVA106 and JAG8810 as tolerant cultivars, and Clodano, DRK7024 and M82 as sensitive cultivars.

It has been described that under NS conditions, tomato floral bud size is generally synchronized with the developmental stage of the reproductive organs (Brukhin *et al*., 2003; **Figure 3 A-D**). Therefore, we determined first, under our NS growth conditions, the developmental stage of the male gametophyte in relation to floral bud size, using a histological approach. Four pollen developmental stages were considered because they were easily recognizable: microspore mother cell (**Figure 3A**), present in 4.5 mm buds in WVA106; Meiosis (**Figure 3B**), in 5.5 mm buds in WVA106; Tetrad (**Figure 3C**) in 6 mm buds in WVA106 and Mature pollen grain (**Figure 3D**) in 7mm buds in WVA106. Most of the time, only one pollen developmental stage was observed for a determined bud size under NS conditions, indicating the synchronization of floral bud and pollen development in the different cultivars studied.

**Figure 3.**
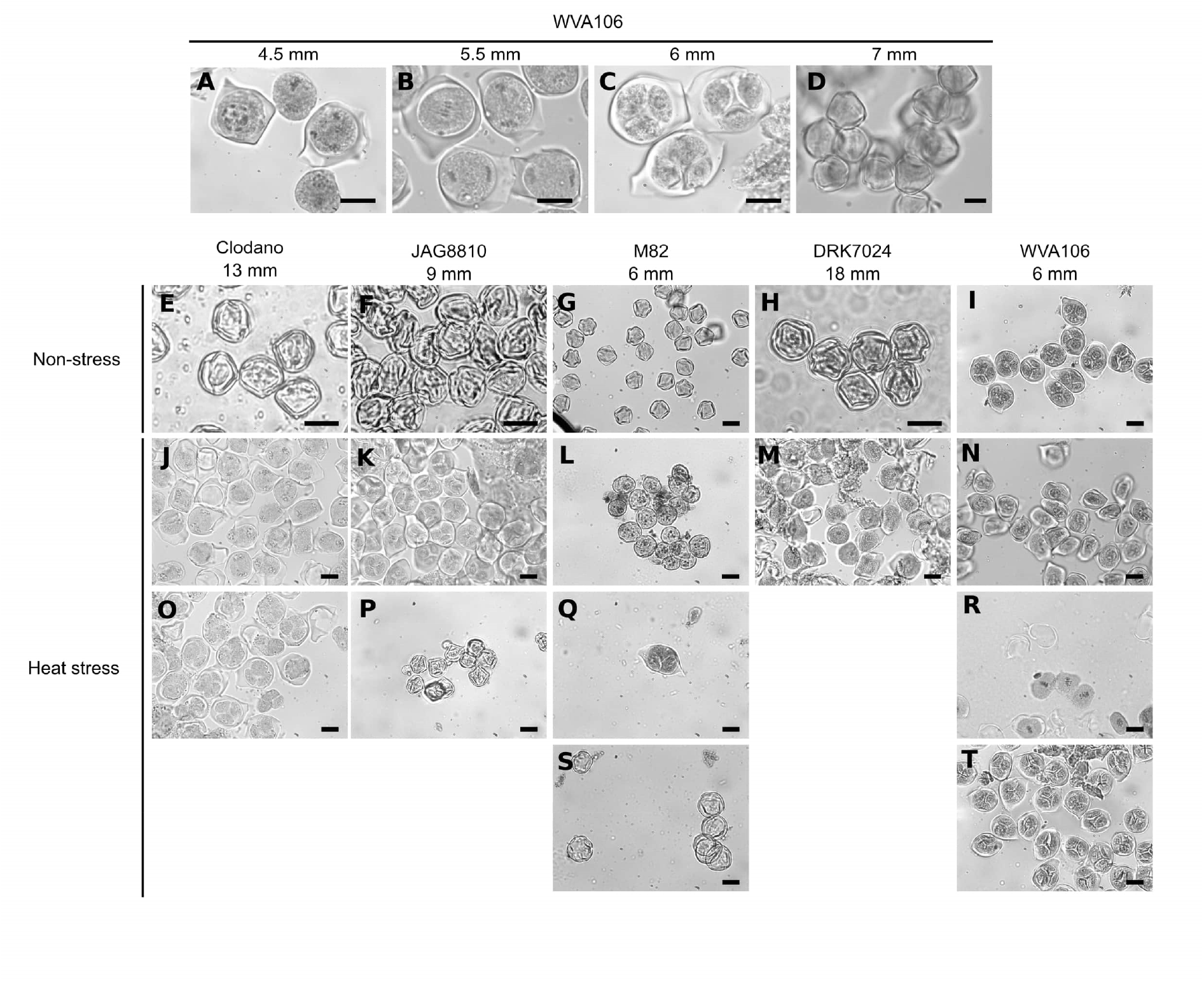
Histological analyses of pollen development in relation with bud size under non-stress (NS) and heat stress (HS) conditions in different tomato cultivars. Pollen developmental stages observed are the following: microspore mother cell (A, L, M, N); Meiosis (B, J, R); Tetrad (C, I, K, O, Q, T); Mature pollen grains (D, E, F, G, H, P, S). Bar, 20µm.

Indeed, Clodano, JAG8810, M82 and DRK7024 floral buds at 13mm, 9mm, 6mm, and 18mm respectively only contained mature pollen grains (**Figure 3 E-H**) and 6mm floral buds in WVA106 only contained tetrads (**Figure 3 I**). Under HS conditions, all cultivars, except DRK7024, showed several pollen developmental stages for a determined bud size (**Figure 3 J-T**). For DRK7024, while only one pollen developmental stage was observed at a time, pollen development was delayed compared to NS condition as indicated by the observed pollen mother cells instead of mature pollen in 18mm floral buds (**Figure 3 H and M**).

These results were confirmed when reporting the size of the flower buds in relation to pollen developmental stage. In three cultivars (Clodano, DRK7024 and JAG8810), the stressed flower buds were longer and showed more variability than in NS conditions for most pollen developmental stages, thus suggesting a delay in pollen development (**Figure 4**). Conversely, floral bud size for a defined pollen developmental stage in WVA106 and M82 was not different indicating that pollen development was not delayed but rather desynchronized under HS condition (**Figure 4**). Altogether these results indicate that pollen development is either delayed (Clodano, DRK7024) and/or desynchronized (Clodano, JAG8810, M82, WVA106) by HS, regardless of the observed tolerance to the heat wave in terms of pollen germination capacity as determined in Figure 2.

**Figure 4.**
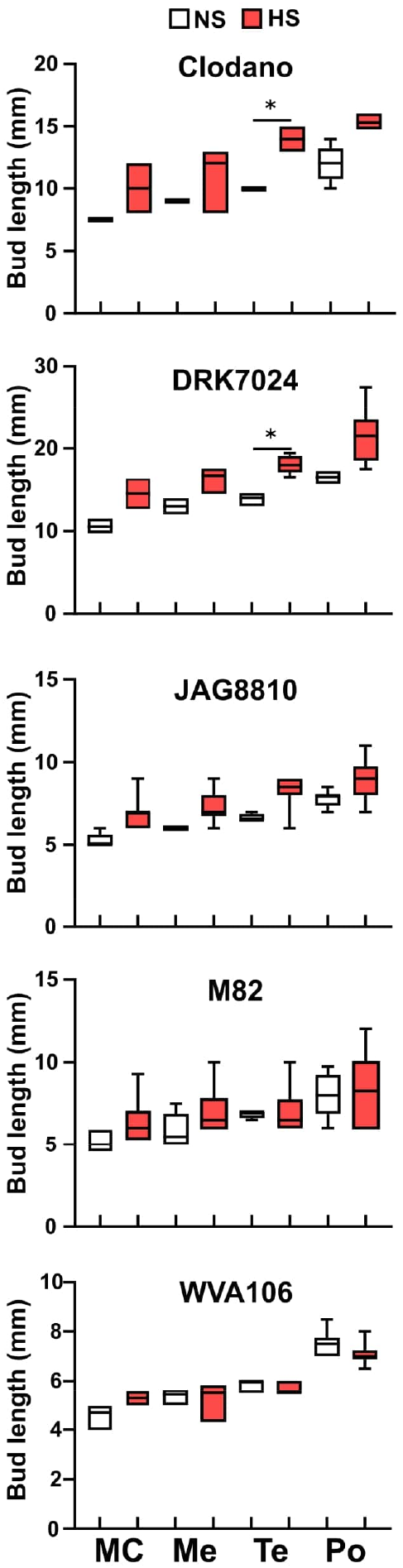
Size of floral buds for a defined pollen developmental stage in Clodano, DRK7024, JAG8810, M82, WVA106,). Pollen Mother Cell (MC); Meiosis (Me); Tetrad (Te); Mature Pollen (Po). * p-value<0.05 (Kruskal Wallis test).

### Genome-wide expression profiling in response to elevated ambient temperatures

In an attempt to identify the molecular pathways differentially regulated in the five selected tomato cultivars displaying contrasting behaviors for pollen germination and floral development under elevated temperature, we carried out a genome-wide transcriptomic analysis by RNAseq. RNA was extracted from floral buds of each tomato cultivar grown under NS or HS conditions (3 weeks at 35°C during the day (16h) and 25°C during the night (8h)). Based on the previous histological analysis, floral buds were harvested at the tetrad stage (tetrad floral bud, TFB) and at the pollen mature stage (pollen floral bud, PFB) based on their size **(Table S1)**.

Following quality check and pre-processing of the sequence data, the reads were mapped against the SLmic1.0 tomato reference genome (gene model version 1.1). Principal Component Analysis (PCA) of the RNA sequencing data showed that the samples clustered together according to both developmental stages and growth conditions factors explaining respectively about 60% and 10% of the whole variance (**Figure S3**). In TFB and PFB, the transcriptomic profiles corresponding to samples subjected to HS were clearly separated from the ones in NS conditions, indicating that the simulated heat wave triggered a transcriptional response in all cultivars.

### General and common transcriptomic response to HS

Differentially expressed genes (DEGs) in HS versus NS conditions were computed for each condition and in the two developmental stages as described in material and methods. The RNA-seq analysis yielded a total of 26017 expressed genes among which 3647 were significantly differentially expressed (adjusted p-value<0.05, 1<Log2FC< −1) across the five cultivars at the two stages and between the two growth conditions (**Table S2**). To analyze the global response to HS at each developmental stage independently of the cultivar considered, the down- or up-regulated genes from all cultivars were pooled and classified based on their GO annotation focusing on biological processes. In both PFB (1415 up and 1420 down, **Table S3**) and TFB (1369 up and 929 down, **Table S3**), most DEGs having enhanced or decreased expression in response to HS belonged to GO classes linked to response to stimulus, response to stress and response to abiotic stimulus (GO:0050896; GO:0006950 and GO:0009628 respectively) corresponding to 20 to 40% of the set of genes with an associated GO term in the studied list **(Table S4)**. As expected, the categories “response to heat” (GO:0009408) and “cellular response to heat” (GO:0034605) were found enriched in the upregulated DEGs. Additionally, “response to reactive oxygen species” (ROS, GO:0000302), “response to oxygen-containing compound” (GO:1901700) and “cellular response to oxygen-containing compound” (GO:1901701) were enriched categories, containing around 15% of the PFB and TFB up-regulated genes. Interestingly down-regulated genes corresponding to the GO “sporopollenin biosynthetic process” (GO:0080110) were found enriched in PFB suggesting that pollen development might be affected. This global analysis thus reveals that the floral buds at both tetrad and pollen developmental stages were severely affected by the heat wave resulting in an oxidative stress.

We then analyzed the DEGs commonly up- or down-regulated at PFB and TFB stages. 820 out of 1964 (42%) and 534 out of 1815 (29%) DEGs were found to be commonly up-regulated and down-regulated, respectively in PFB and TFB samples, highlighting the response similarity to HS independently of the developmental stage (**Figure 5A-B**). Most DEGs with enhanced expression in response to HS belonged to GO classes linked to response to stimulus such as response to heat (GO:0009408, GO:0034605), response to ROS (GO:0000302 and GO:1901700), and cellular processes such as protein folding (GO:0006457 and GO:0042026) (**Figure 5C; Table S5)**. For the down-regulated DEGs the GO enrichment analysis revealed mainly classes related to metabolic processes such as photosynthesis (GO:0009765). These results indicated that the global tomato transcriptome response at both TFB and PFB stages is largely common and suggested that both tissues undergo inhibition of photosynthetic processes.

**Figure 5.**
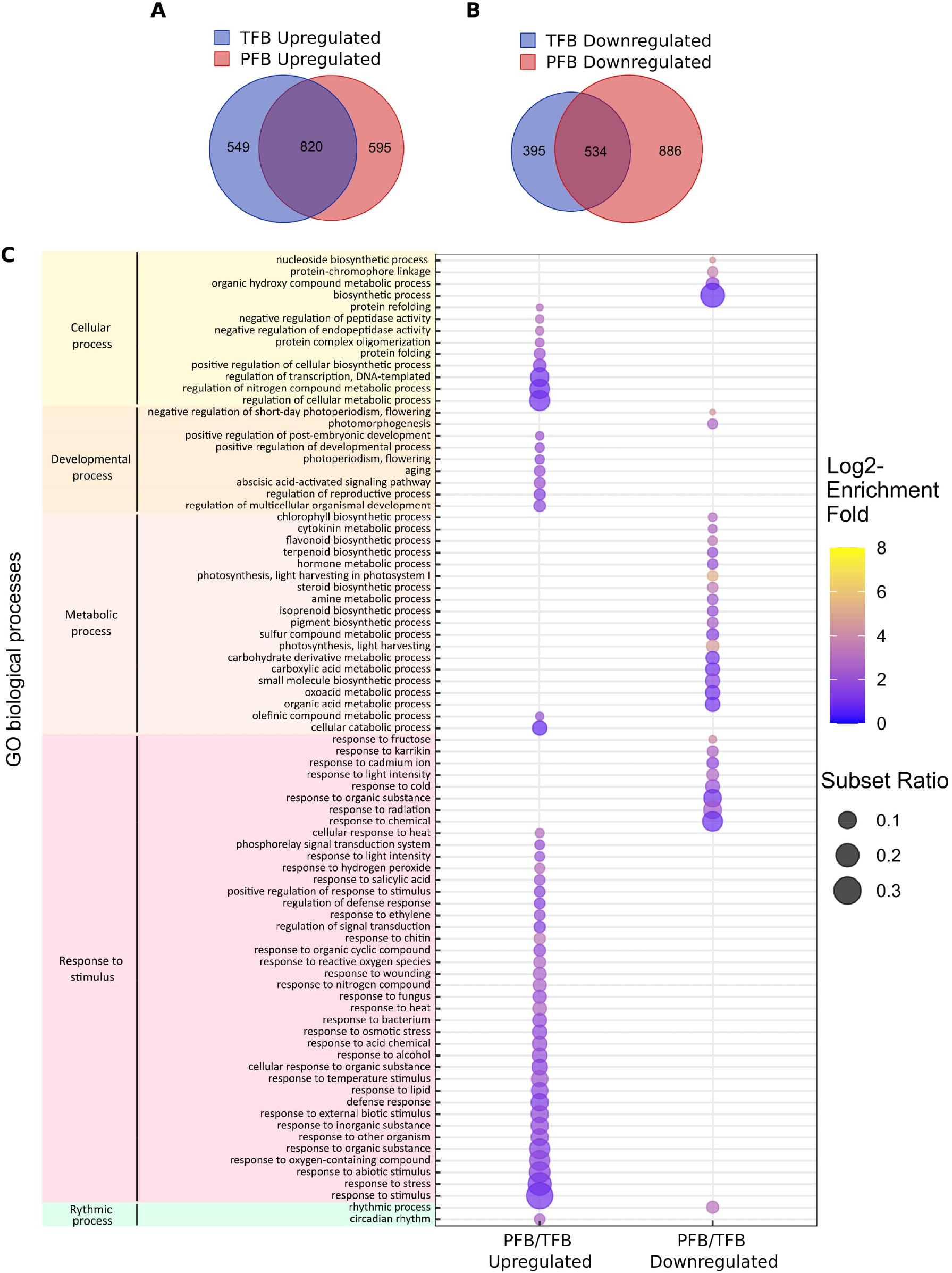
Global transcriptome response to HS in all tomato cultivars. (A-B) Venn diagram presenting the overlap of DEGs between upregulated (A) and downregulated (B) genes between TFB and PFB stages in all cultivars. (C) Gene ontology (GO) enrichment analyses of the common DEGs between PFB and TFB. The dot size is representative of the number of DEGs associated with the process and the fold enrichment is according to a heat map (dot color).

### Cultivar specific transcriptomic response to HS

To evaluate the common and specific responses to HS between cultivars, we first compared the number of DEGs in the different tomato cultivars under HS condition. We noticed that this number was higher in the pollen sensitive-than in the pollen tolerant cultivars: at the TFB stage, we found 872 and 742 DEGs for JAG8810 and WVA106 respectively, as the two tolerant cultivars, and 1233, 1089 and 1293 DEGs for M82, DRK7024 and Clodano respectively, as the three sensitive cultivars (**Table S3**). At the PFB stage, 587 and 1402 DEGs were found for JAG8810 and WVA106 respectively, and 1327, 1418 and 1388 for M82, DRK7024 and Clodano respectively. These observations suggested that the higher number of responding genes could be indicative of a higher sensitivity of pollen to the HS.

We then compared the DEGs found in the five cultivars at each developmental stage. The Venn diagram of the up-regulated genes in TFB showed that 147 genes were common between all cultivars which represent between 17% (for Clodano) and 32% (for WVA106) of the up-regulated DEGs (**Figure 6A, Table S6)**. In PFB, 57 genes were found common between the five cultivars representing between 7% (for M82) and 15% (for JAG8810) of the up-regulated DEGs (**Figure 5B, Table S6**). Interestingly only 13 out of 502 genes were found specifically up-regulated in JAG8810 indicating that the response at this stage in this cultivar is almost completely shared by the others. Concerning the down-regulated DEGs, 97 genes (representing between 20% for DRK7024 and 34% for WVA106) and 69 genes (representing between 9% for DRK7024 and 32% for JAG8810) were commonly down-regulated in TFB and PFB respectively between all cultivars (**Figure 5C and 5D, Table S6**).

**Figure 6:**
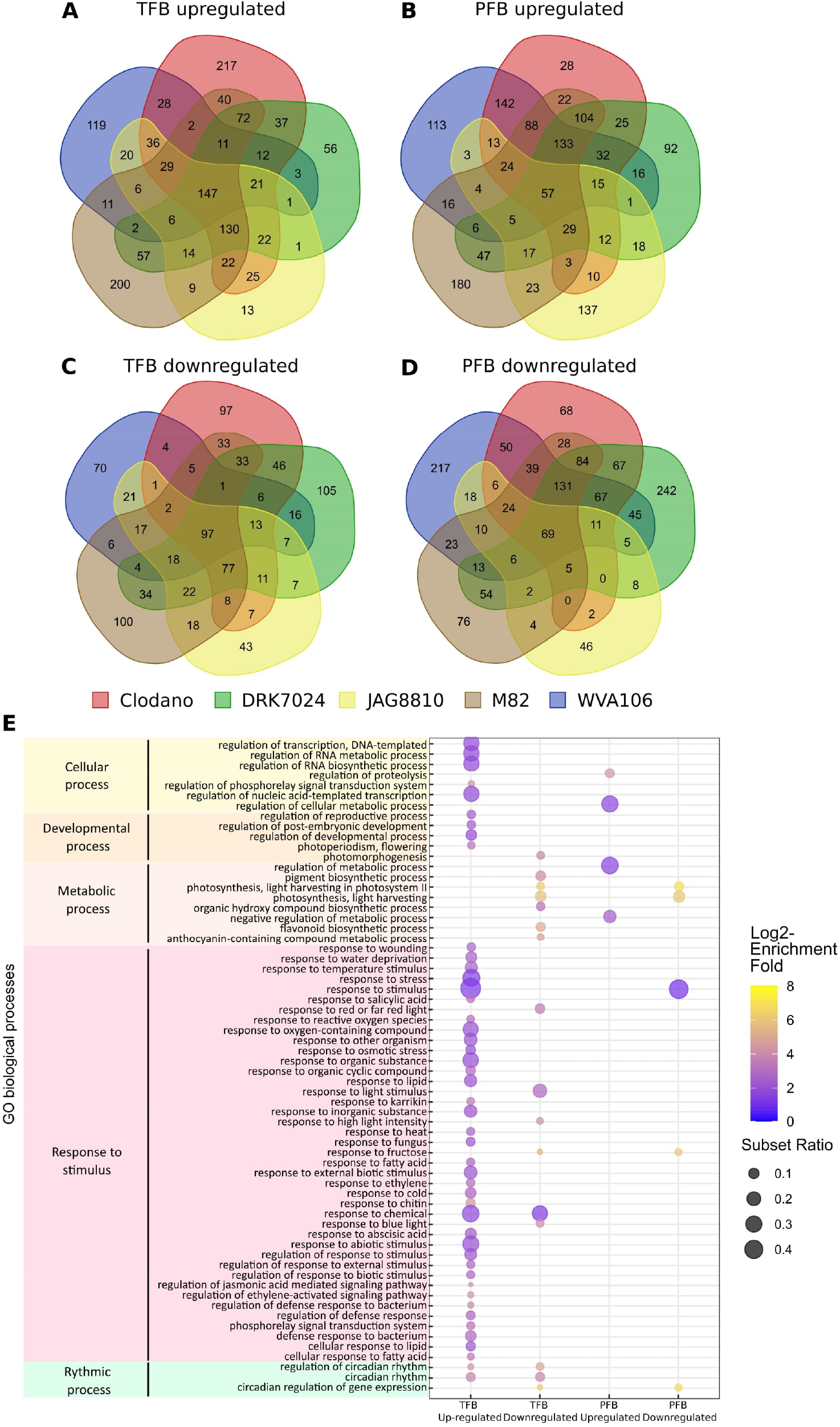
Common transcriptome response to HS between all cultivars (A-D) Venn diagram presenting the overlap of DEGs between all cultivars: upregulated genes in TFB (A) and in PFB (B); downregulated genes in TFB (C) and in PFB (D). (E) Gene ontology (GO) enrichment analyses of the common DEGs between all cultivars. The dot size is representative of the number of DEGs associated with the process and the fold enrichment is according to a heat map (dot color).

The 147 commonly up-regulated DEGs identified in TFB were mainly involved in response to stimulus (GO:0050896) with 49% of the set of the genes with an associated GO term in the studied list (**Figure 5E and Table S7**). As expected the GO term “response to heat” (GO:0009408) and “response to oxygen-containing compound” (GO:1901700) were found enriched. Similarly, GO terms corresponding to hormonal responses such as salicylic acid (GO:0009751) and ethylene (GO:0009723), as well as regulation of reproductive process (GO:2000241) were also enriched. At PFB stage, GO terms corresponding to metabolic process (GO:0019222) were found enriched among the 57 commonly up-regulated DEGs. The GO enrichment of the down-regulated DEGs at both TFB and PFB stages revealed mainly photosynthesis related genes (GO:0009765) indicating that both stages are affected in this process during HS. Altogether these results suggested that the common response to HS in the five cultivars mainly corresponds to the upregulation of heat stress response genes, oxidative stress and hormonal pathways and the down-regulation of photosynthesis related genes.

To gain some insight into the common and specific responses of the two developmental stages studied, we then compared the DEGs between PFB and TFB in each cultivar. Interestingly, the percentage of commonly up-regulated DEGs between PFB and TFB was higher in sensitive cultivars (36.5% in Clodano; 42.6% in DRK7024 and 43.3% in M82) than in tolerant cultivars (17.5% in WVA106 and 17.0% in JAG8810) (**Figure S4**). A similar result was observed for commonly down-regulated DEGs with 22.7% in Clodano, 32.5% in M82 and 31% in DRK7024 compared with 16.0% and 17.5% in WVA106 and JAG8810 respectively (**Figure S4**). This might be the consequence of the observed developmental alterations and desynchronization of pollen development in sensitive cultivars in which PFB still contain non-mature pollen that might share more DEGs with TFB compared to pollen of tolerant cultivars in which the pollen is fully mature in PFB.

### Flower, pollen and anther development related genes mis-expressed upon HS

The “regulation of reproductive process” (TFB and PFB; GO:2000241; 44 genes) and “developmental process involved in reproduction” (PFB, GO:000300; 130 genes) categories were found enriched in the pool of up-regulated and down-regulated DEGs for all cultivars (**Table S8**), confirming that HS affected flower development. Given the fact that the applied HS affects pollen development, we further examined the transcriptional expression of several genes expressed during tomato pollen development (Chaturvedi *et al*., 2013) under NS and HS conditions. We also included in our dataset genes proposed to be related to tomato pollen- and tapetum development (Jeong *et al*., 2014; Liu *et al*., 2019) (**Table S9**). Few genes specifically expressed during anther development and in the microspore were found to be down-regulated under HS in several cultivars, although no clear response pattern was observed (**Table S9**). Interestingly, under NS conditions, genes encoding proteins displaying high expression levels in pollen mother cells such as histones (histone H4, Solyc04g011390; histone H2A, Solyc09g010400), proteasome α-subunit (Solyc07g055080) and ribosomal proteins (S17, Solyc11g006690 and L12, Solyc06g075180) displayed higher expression in TFB than in PFB in NS conditions. However, the fact that these genes are not up-regulated under HS in PFB samples suggests that the observed cytological desynchronization of pollen development is not visible at the gene expression level. Conversely, genes encoding proteins specifically expressed in mature pollen (Jeong *et al*., 2014) such as the anther-specific protein TA29 (Solyc02g078370), TomA108 (Solyc01g009590), a gene specifically expressed in anthers and tapetum (Xu, Liu and Chen, 2006), a cysteine protease (Solyc07g053460) or AtAMS-like (Solyc08g062780), encoding a fatty acyl-CoA reductase important for pollen wall development (Chen *et al*., 2011) were specifically expressed in PFB in NS condition. While validating the sampling stages chosen based on cytological observations, these observations did not reveal any striking differences between NS and HS conditions concerning pollen-related genes.

### Genes encoding HSPs and HSFs are mis-expressed under HS

Among the genes up- and down-regulated, we identified a high number of *HSPs* (heat shock proteins) encoding chaperone proteins and *HSFs* (heat shock factors) encoding DNA-binding proteins, known to be HS related genes. 48 and 44 out of 208 tomato HS-related genes (Keller *et al*., 2018) were found up-regulated in at least one cultivar in TFB and PFB respectively, for a total number of 50 genes at both stages (**Table S10**). Interestingly, more *HSPs* were found upregulated in sensitive (45 for Clodano; 33 for DRK7024; 22 for M82) than in tolerant (20 for WVA106 and 20 for JAG8810) cultivars. Moreover, for the 13 commonly up-regulated *HSPs* in all cultivars, despite a similar absolute expression level under NS conditions, the mean log2-fold change between NS and HS conditions in sensitive cultivars was higher than in tolerant cultivars for the two stages analyzed (2.7-3.8 for Clodano; 2.8-2.7 for DRK7024; 2.4-2.8 for M82; 1.4-1.7 for WVA106 and 1.8-1.5 for JAG8810 (values indicated for TFB-PFB)) suggesting that the sensitive cultivars might sense the stress in a stronger manner. Among these 13 commons genes, 11 encode transcription factors belonging to the HSF class, and 2 to the protein folding class HSP40. Several of these genes have been already described as important factors for HS tolerance in male reproductive tissues and particularly at the tetrad stage such as HsfA2, HsfA6b (Fragkostefanakis *et al*., 2016). These results indicate that depending on the cultivar considered, the level and diversity of HS related proteins induced may vary, suggesting that this differential behavior could lead to the differences observed in the responses to HS.

### ROS related genes mis-expressed under HS

A total of 259 genes (out of 1860) related to the response to oxygen-containing compound GO (GO:1901700) and 89 genes related to the response to oxidative stress (GO:0006979) were up-regulated at both stages in all cultivars (**Table S11**). Genes encoding ROS scavenging enzymes such as the Ascorbate peroxidase (Solyc09g007270), already reported to be responding to HS in tomato microspores (Frank *et al*., 2009), was found upregulated in HS condition. Antioxidant enzymes such as Catalases (Solyc04g082460 and Solyc12g094620) were upregulated under HS and Solyc12g094620 displayed higher expression levels under NS condition in tolerant cultivars compared to sensible cultivars in both TFB and PFB stages (between 2.1- and 2.6-fold increase in TFB and between 1- and 2-fold in PFB). Moreover, enzymes involved in ROS detoxication such as the iron superoxide dismutase (Solyc06g048410) were found up-regulated.

## Discussion

### Various tomato cultivars respond unequally to HS

The effects of different types of stresses, including HS, on flower development have been studied for a long time. It is known that the optimal temperature for growing tomato plants is 25°C during the day and 20°C at night (Alsamir *et al*., 2017). For most tomato cultivars, if the temperatures exceed 26ºC during the day and 20ºC during the night, fruit set is impaired mainly due to floral development alteration, and therefore fruit yield is reduced (Sherzod *et al*., 2020). Because the response to stress varies widely between genotypes (Arena *et al*., 2020; Gonzalo *et al*., 2021), gaining deeper knowledge of the developmental and molecular response of various cultivars may provide clues for innovative breeding strategies aiming at improving commercial tomato cultivars for better adaptability to climate change. In the present work, we investigated the responses to heat stress in eleven tomato cultivars producing fruits of different sizes and shapes. The data show that one of the most characteristic effects of HS on floral morphology is stigma exertion, which negatively impacts self-pollination due to the excessive elongation of the styles minimizing pollen access to the stigmas and reducing fertilization, and consequently decreasing fruit yield and fruit quality (Fernandez-Muñoz and Cuartero, 1991; Giorno *et al*., 2013). A similar response was previously described in the tomato cultivar Micro-Tom (Pan *et al*., 2019) and the wild species *S. pimpinellifolium, S. pennellii* and *S. chillense* (Alsamir *et al*., 2017) as a consequence of increased temperature. Interestingly, we only observed this effect in the cultivar Sassari. The absence of stigma exertion in the other cultivars could be explained by the milder stress applied in our study compared to the harsh heat condition used in previous studies (35°C/30°C and 16 h/8 h day/night) for 12 days in Pan *et al*. (2019) and two months in a tunnel house with max of 50ºC and min of 30ºC in Alsamir *et al*. (2017). The pistil exertion in the Sassari cultivar could thus indicate that this cultivar is very sensitive to HS. This high sensitivity is also highlighted by the absence of pollen germination under the HS conditions we applied (**Figure 2**). As previously observed (Giorno *et al*., 2013), HS triggered additional morphological alterations including curled petals and sepals, and early opening of the petals that were more or less pronounced depending on the cultivar. The morphological observations after HS thus reveal that the cultivars respond differently to the stress applied.

### Under HS, pollen germination was affected in all cultivars but with different ranges

Pollen is easily damaged by exposure to HS (Bita *et al*., 2011; Paupière *et al*., 2017; Rieu, Twell and Firon, 2017). The exposure of tomato plants to high temperatures has two effects on pollen grains: a reduction in the number of grains and a reduction in germination and viability (Pressman, Peet and Pharr, 2002). The impact of HS on pollen germination was described previously in many species such as in rice (Coast *et al*., 2016), sorghum (Sunoj *et al*., 2017), peanuts (Kakani *et al*., 2002), spring wheat (Bheemanahalli *et al*., 2019) and tomato (Sherzod *et al*., 2020). Thermotolerant tomato cultivars have been defined as plants with higher yield, high quantity of viable pollen grains, and higher germination capacity under HS conditions when compared to thermosensitive plants (Firon *et al*., 2006). In our study, we found that two cultivars were possibly thermotolerant, namely WVA106 and JAG8810, since they maintain high pollen germination under HS. However, these two cultivars differ greatly with regard to their germination capacity under NS conditions (82.7% and 23.4%, respectively). These variations were also observed in other studies as in spring wheat (*Triticum aestivum*) where pollen germination of 22 cultivars was compared and a germination percentages of 87% maximum and 30% minimum were observed (Bheemanahalli *et al*., 2019). The decrease in the capacity of pollen to germinate under HS conditions is probably due to damages in the physical structure of pollen, pollen wall composition or lower energy (sugar) status (Jiang *et al*., 2015; Sita *et al*., 2017; Sunoj *et al*., 2017; Djanaguiraman *et al*., 2018; Jagadish *et al*., 2021). JAG8810 might be more thermotolerant than WVA106 since the decrease of pollen germination was less important than for WVA106. In other cultivars, we found that pollen germination was lower than 10%, a very low rate probably leading to reduced fruit set. These findings indicate that these 9 cultivars are sensitive to the stress applied when considering the pollen germination trait. For that trait too, it seems that the genetic background influences differently the response to HS.

### HS affects pollen development in the sensitive cultivars

Pollen grains are formed inside the anthers through a series of developmental steps from microsporocyte meiosis to pollen release (Gómez *et al*., 2015). In general, pollen development is synchronized until microsporocyte meiosis I, and sometimes until meiosis II (Brukhin *et al*., 2003). Cell division synchronization is necessary for successful and coordinated pollination and fertilization (Magnard *et al*., 2001). Specifically, an environmental stress (extreme temperature or drought) induces asynchrony during pollen development and gives rise to meiotic abnormalities (Pécrix *et al*., 2011; De Storme and Geelen, 2020). When cultivated under continuous mild heat, a simultaneous reduction in pollen viability and appearance of anthers with pistil-like structures was observed in tomato indicating that HS impairs anther development (Müller *et al*., 2016). In our study we show that pollen development is affected at very early stages by HS, impairing meiosis and leading to a delay or a desynchronization of pollen development depending on the cultivar considered. At the molecular level, few genes related to pollen development were mis-regulated under HS, although no clear response pattern was found between cultivars or time points.

### Stress-related genes are commonly induced in the various cultivars

Under a short HS treatment, the *HSP* genes are strongly induced and play important roles to protect the plant. HSP act to maintain the homeostasis of the plant as they have a chaperone activity to limit the misfolding of proteins, but also to protect protein complexes like photosystem II (Vierling, 1991; Tsvetkova *et al*., 2002; Waters, 2013). Our global transcriptome analysis showed that all cultivars respond to heat wave by maintaining an increased expression of numerous *HSF* and *HSP* genes and a stronger upregulation in sensitive than in tolerant cultivars. As indicated in previous studies (Fragkostefanakis *et al*., 2016) these results not only strongly suggest that pollen tolerance to HS is linked to the ability of the plant to mitigate the HS effects through a higher expression of heat-response factors, but also by activating a common response for short term extreme HS and heat wave exposure to moderate high temperatures.

## Conclusion

Our results showed that heat waves are detrimental for pollen development in all tested tomato cultivars but the tolerance to heat stress widely varies between genotypes. The effects of global warming with the increase in heat waves magnitude and frequency on reproductive organs development need to be investigated using different approaches than those implemented for the study of extreme short high temperature-stress. The transcriptome dataset generated in the present study revealed the complex response of tomato to heat and the common and specific response of each tomato cultivar.

## Experimental procedures

### Plant material and growth

Plant material consisted in eleven different tomato (*Solanum lycopersicum*) cultivars: Brioso (Rijk Zwaan), Clodano (Syngenta™), Docet (Monsanto™), DRK7024 (De Ruiter™), JAG8810 (Bayer™), M82, Marbonne (Gautier Semences), Moneymaker, Rebelski (De Ruiter™), Sassari (Rijk Zwaan) and West Virginia 106 (WVA106). In NS conditions, plants were cultivated in a greenhouse with a photoperiod of 16h/8h with a mean temperature of 24°C during the day and a mean temperature of 18°C during the night. For heat-stress treatments, the plants at bolting stage have been submitted during 3 weeks to 35°C during the day (16h) and 25°C during the night (8h). In both conditions, the hygrometry was around 55% all day and plants had a daily watering.

### Pollen developmental stage analysis

To analyze pollen development, floral buds of different sizes were dipped in Carnoy’s fixative solution (60% (v/v) ethanol, 30% (v/v) chloroform, 10% (v/v) acetic acid). Then, the anthers from the flower buds were dissected in acetocarmine staining solution (1% (p/v) carmine 40, 0,5% (p/v) ferric chloride, 45% (v/v) glacial acetic acid) using gauge needles under a stereomicroscope and squashed between a slide glass and a coversleep to release the male reproductive cells as described by (Puchtler *et al*. 1968). Pollen developmental stages were observed using a bright field microscope (Zeiss, Axioplan) and photographed with a CCD camera (Motic 3 megapixels).

### Pollen germination assays

Pollen grains from tomato flowers at anthesis stage were sprayed on the top of pollen germination medium (18% sucrose, 0.01% Boric acid, 1mM CaCl_2_, 1mM Ca(NO_3_)_2_, 1mM MgSO_4_, 0.5% agar; pH=7). After 16 hours of incubation at 25°C in the dark, the preparation was observed using a stereomicroscope Olympus SXZ16 and photographed using a camera (Motic 10 megapixels). The pollen germination rate expressed as a percentage was determined by dividing the number of germinated pollens corresponding to those emitting a pollen tube by the total number of pollen grains.

### Genome wide expression profiling

Total RNAs were isolated from 200 and 500 mg of floral buds at different developmental stages using TRIzol Reagent (Life Technologies). Total RNA extract was purified with the Qiagen RNAeasy mini kit RNA. After DNase treatment (DNA-free Kit, Life Technologies), the total RNA quantity and quality (RNA integrity number, RIN) were evaluated using an Agilent 2100 Bioanalyzer (Agilent Technologies). Only RNA extracts with a RIN of 10 were used for sequencing. The RNA libraries were constructed as described in the Illumina TruSeq Stranded mRNA guide. mRNAs were sequenced in a HiSeq 3000 sequencing system with 2 × 125 bp paired-end sequences (Illumina HiSeq SBS Kit v4) by the Genotoul bioinformatics platform, Toulouse (http://bioinfo.genotoul.fr/index.php).

Three biological replicates per plant and per stage were harvested for NS and heat-stress conditions, resulting in 60 samples (5 cultivars × 2 conditions × 2 developmental stages × 3 biological replicates). To obtain the genome-wide expression profiling, RNA samples were subjected to next-generation sequencing.

For each RNA sample, more than 20 million of paired-end reads (10 million fragments) were generated, using an Illumina Hiseq 3000 platform. More than 95% of total clean reads were mapped against the SL3.0 version of the reference tomato genome for read mapping (ftp://ftp.solgenomics.net/genomes/Solanum_lycopersicum/Heinz1706/assembly/build_3.00/) and almost 90% of the mapped reads matched a feature on the related gene model.

### Bioinformatic analyses

Statistical analyses and graphs have been performed with the R software and homemade scripts. The DE analysis has been carried out with the DESeq2 R-package (Love *et al*., 2010). Some analyses, such as the PCA analysis, were carried out with functions contained in the DESeq2 package. Reads were checked using fastQC, cleaned using trimGalore, and mapped using Star, a spliced aware mapper software on the new tomato genome SLmic1.0 generated in the frame of TOMGEM project (http://tomatogenome.gbfwebtools.fr/). The mapping was guided using the gene model annotation version 1.1. FeatureCount was then used to calculate the read counts for each gene from each mapping file. A normalization step was performed in order to obtain comparable expression values between conditions and between genes. For this purpose, our pipeline takes into account the relative size of studied transcriptomes, the library sizes and the gene lengths, as described in Maza *et al*. (2013) and Maza (2016). After data normalization using DESeq2 package, a PCA analysis has been conducted with expression data in all conditions and replicates, in order to check global sample variability.

Genome-wide expression data have been analyzed in order to identify genes differentially expressed between different conditions that have been tested. The “Relative Log Expression” normalization (RLE) implemented in the DESeq2 package has been used as normalization method. To highlight DEGs, the genes exhibiting an adjusted p-value < 0.05 and a log2 fold-change < −1 or > +1 have been selected. To eliminate very lowly expressed genes, the up-regulated DEGs with a count value < 10 in HS condition and the down-regulated DEGs with a count value < 10 in NS condition were filtered out.

To calculate the intersections of DEGs lists within the different conditions, Venn diagrams corresponding to textual outputs were generated by using the Venn diagram tools at http://bioinformatics.psb.ugent.be/webtools/Venn/BAR web site at http://bar.utoronto.ca/ or at https://www.biovenn.nl/index.php (Hulsen *et al*., 2008) for area-proportional Venn diagrams. The TomExpress web site, at http://tomexpress.toulouse.inra.fr/query, was used to extract gene expression data obtained from previous genome-wide expression analyses (Zouine et al., 2017).

The GO enrichment analysis has been carried out on PLAZA 5.0 (Van Bel et al., 2022) using the Plaza workbench, with a significance threshold of 0.05 and without any data filter.

## Authors’ contributions

M.H., C.C., F.D., M.B. and N.G. conceived and supervised the study, obtained funding, and provided resources. M.H., F.D. and N.B. designed the methodology. M.H., F.D., N.B. and A.D. performed the experiments. E.M., M.Z. did the RNA sequencing. M.H., F.D., R.M-P and N.B. analyzed the data. N.G., R.M-P and N.B. wrote the original draft. All authors reviewed and edited the manuscript.

## Acknowledgements

We thank Isabelle Atienza for greenhouse management. Research was supported by the European Union’s Horizon 2020 Research and Innovation Program through the TomGEM project under grant agreement Number 679796. R.M-P was supported by the Conselleria de Innovación, Universidades, Ciencia y Sociedad Digital de la Generalitat Valenciana under the grant APOSTD/2019/001.

## Figure legends

**Figure S1.**
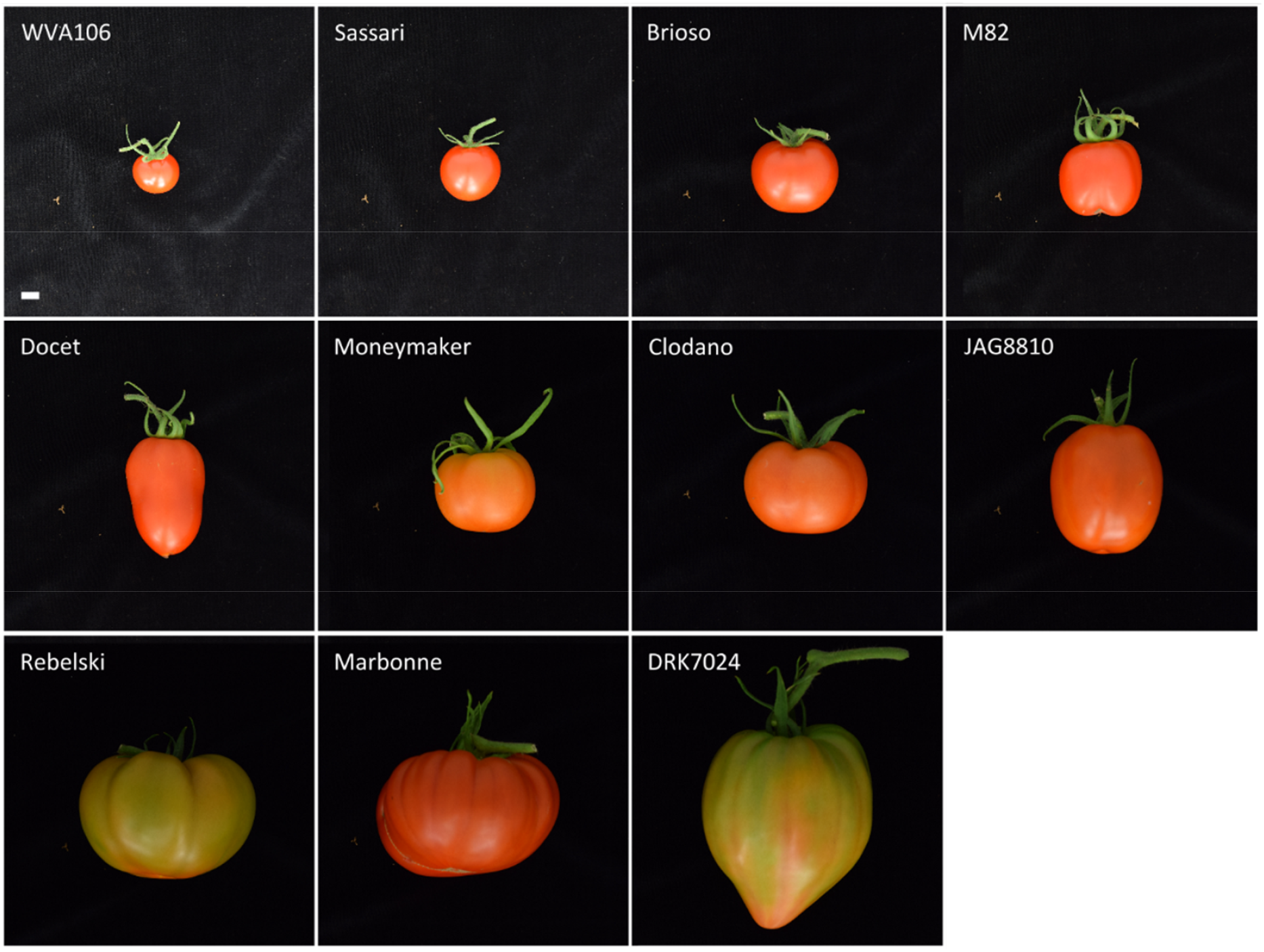
Fruit morphology in the eleven selected cultivars used in the study. Size differences allowed the classification of the 11 cultivars into three main groups: small (WVA106, Sassari, Brioso and M82), medium (Docet, Moneymaker, Clodano and JAG8810), and large fruits (Rebelski, Marbonne and DRK7024). Scale bar = 1 cm (applies to all panels).

**Figure S2.**
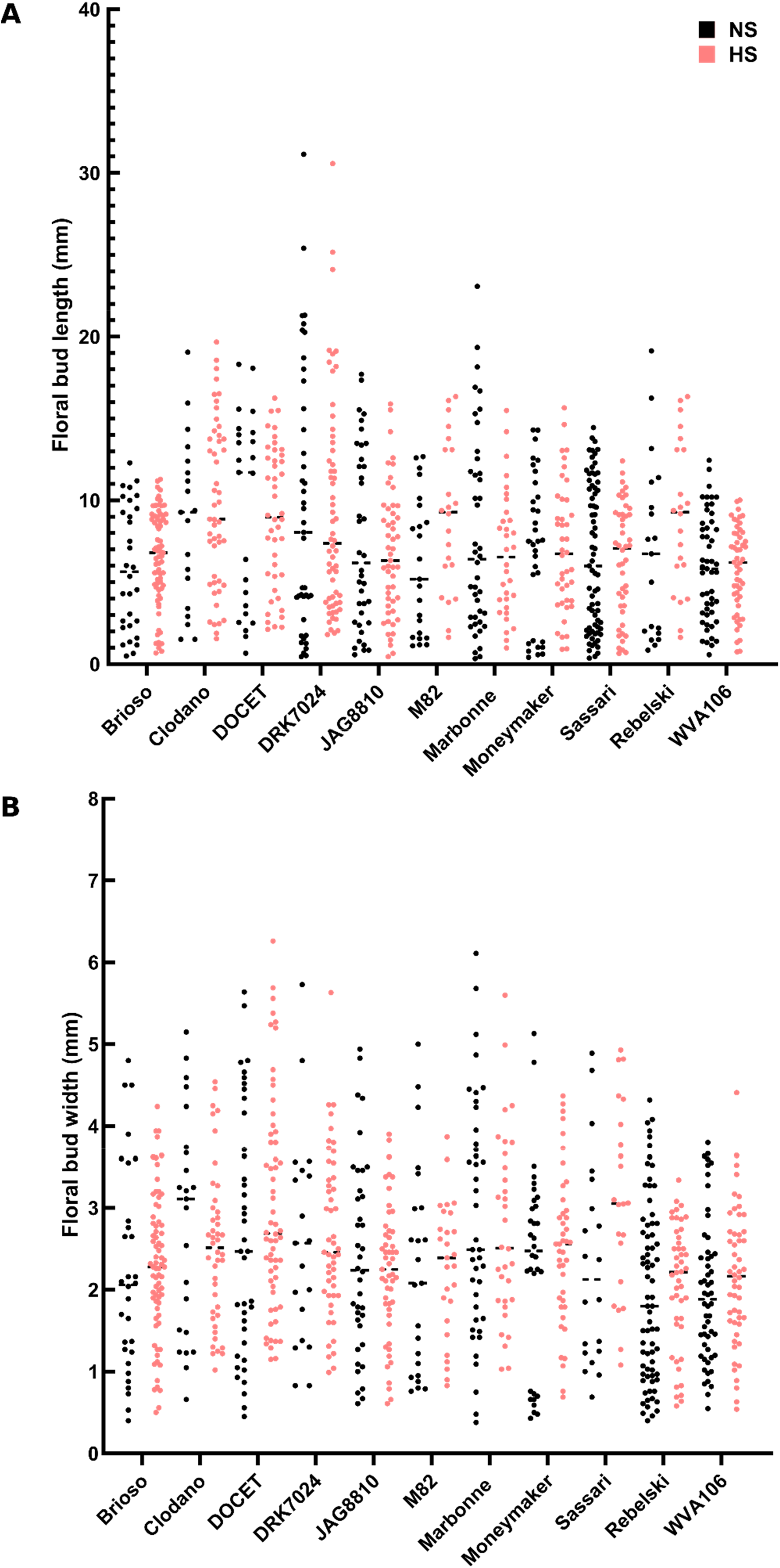
Distribution of floral bud length (A) and width (B) under NS and HS conditions in the 11 cultivars.

**Figure S3.**
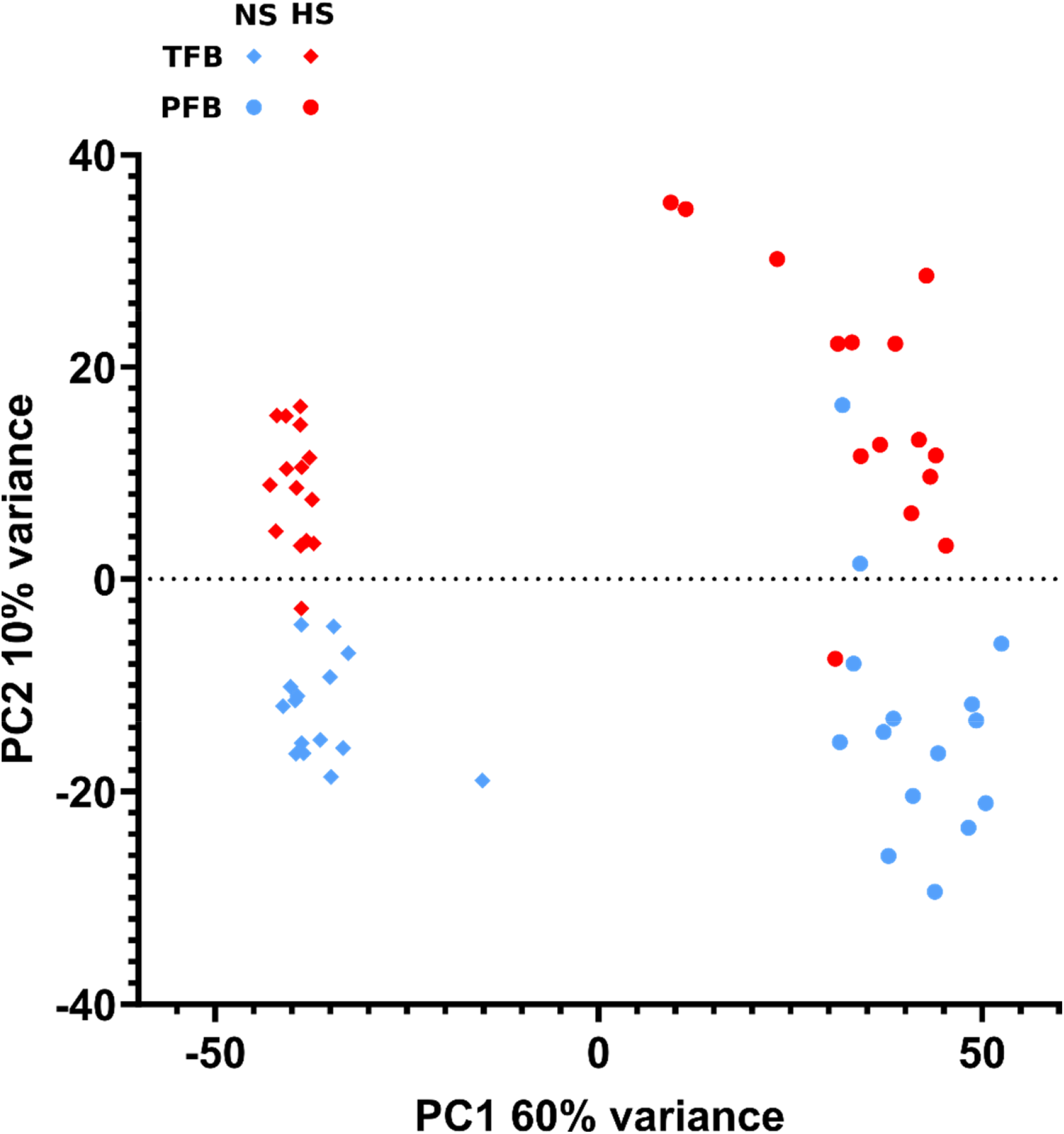
Principal component analysis (PCA) of gene expression in TFB and PFB in NS and HS conditions. PC1, explaining 60% of the total variance, differentiates the TFB stage (diamond) from the PFB stage (circle). PC2, explaining 10% of the total variance, separates the heat stress samples (red) from the control samples (blue).

**Figure S4.**
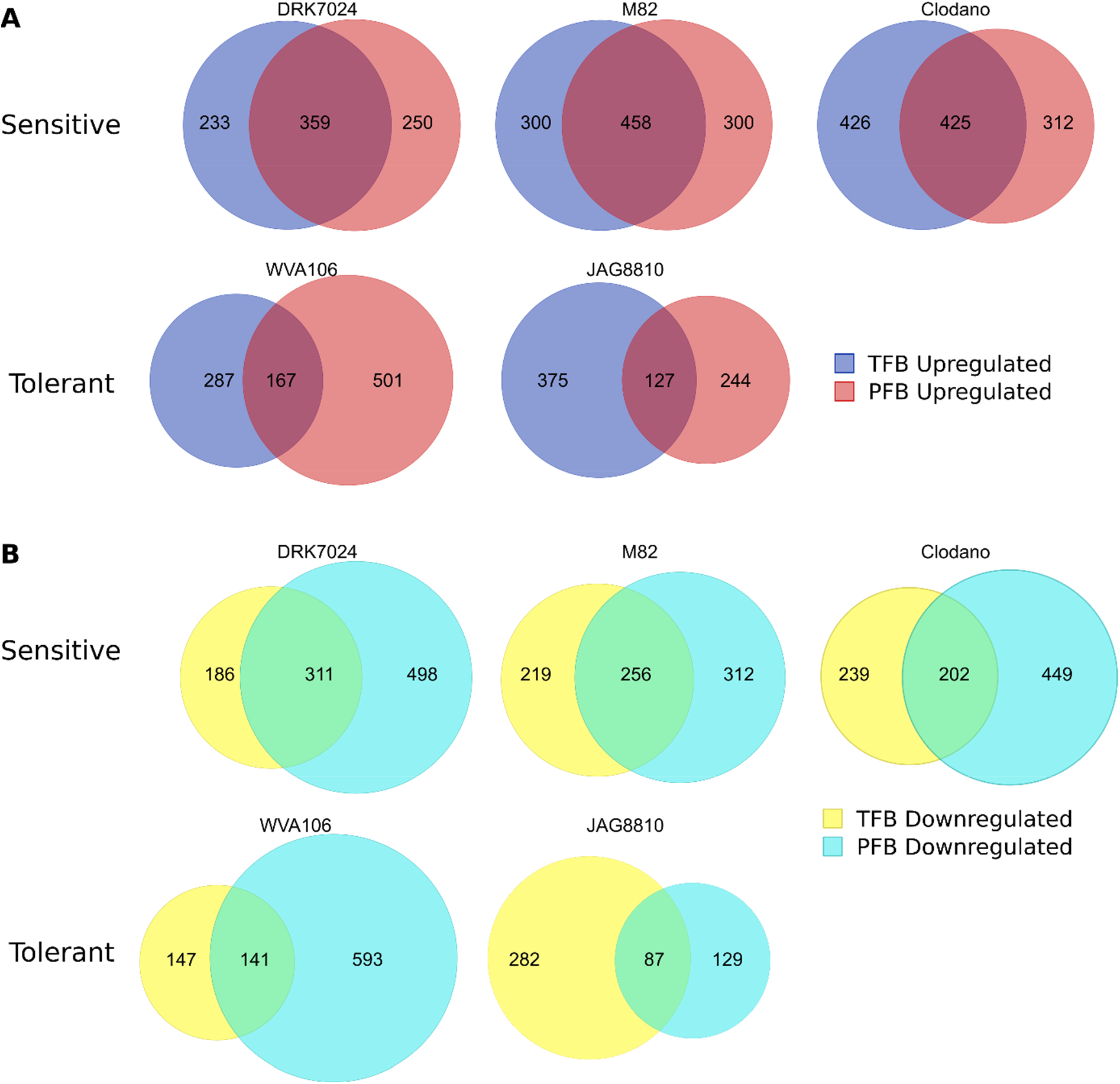
Common and specific responses between PFB and TFB stages in all cultivars Venn diagram analysis of the upregulated (A) and downregulated (B) DEGs between PFB and TFB stages in pollen sensitives and pollen tolerant cultivars

## Tables legends

**Table S1**. Sizes of the floral buds containing tetrads (TFB) and mature pollen grains (PFB) harvested for the transcriptomic analysis.

**Table S2**. List of DEGs under heat-stress conditions in all cultivars in TFB and PFB.

**Table S3**. Number of differentially expressed genes under heat-stress conditions based on a log2 fold-change of 1 and a p-value <0.05.

**Table S4**. Functional categories overrepresented in the list of all up- and down-regulated DEGs across all the tomato cultivars under heat stress.

**Table S5**. Functional categories overrepresented in the list of commonly up- and down-regulated DEGs across all the tomato cultivars in PFB and TFB under heat stress.

**Table S6**. List of common DEGs between all cultivars in TFB and PFB.

**Table S7**. Functional categories overrepresented for each developmental stage in the common DEGs between all cultivars

**Table S8**. (A) List of DEGs corresponding to the GO “regulation of reproductive process” found enriched in the upregulated DEGs of TFB and PFB. (B) List of DEGs corresponding to the GO “developmental process involved in reproduction” found enriched in the downregulated DEGs PFB.

**Table S9**. List of DEGs corresponding to pollen development-related genes and their expression under NS and HS conditions.

**Table S10**. List of DEGs corresponding to HSP and HSF genes and their expression under NS and HS conditions.

**Table S11**. List of DEGs corresponding to GO:1901700 “response to oxygen-containing compound” and their expression under NS and HS conditions.

